# “PDGFRα is required for postnatal cerebral perivascular fibroblast development”

**DOI:** 10.1101/2025.06.07.658411

**Authors:** Hannah E. Jones, Kelsey A. Abrams, Katherine A. Fantauzzo, Julie A. Siegenthaler

**Author notes:** Corresponding Author, Contact Information: Julie Siegenthaler, PhD Associate Professor, University of Colorado, Anschutz Medical Campus Dept of Pediatrics, Section of Developmental Biology 12800 East 19th Ave, MS-8313, Aurora, CO 80045.

## Abstract

Perivascular fibroblasts (PVFs) are a cell type associated with large diameter blood vessels in the brain and spinal cord parenchyma and leptomeninges. PVFs have previously defined roles in injury and neuroinflammatory diseases and predicted roles in supporting neurovascular function. The temporal dynamics of PVF development in the pre- and postnatal cerebral cortex have recently been described, however the molecular mechanisms that underly PVF development have not been identified. PVFs express both platelet derived growth factor receptors (PDGFRs), PDGFRα and PDGFRβ. Here we investigate the role of PDGF signaling in PVF development. We use immunohistochemistry and RNA transcript detection methods to show developmental expression of PDGFRs by PVFs and examine distribution of PDGF ligand expression in the brain. We show that postnatal deletion of PDGFRα in fibroblasts using the *Col1a2-CreERT* mouse line significantly impairs PVF coverage of cerebral vessels at postnatal day 10. Perivascular macrophages, a cell type previously shown to co-develop with PVFs, have impaired cerebral vessel coverage in conditional mutants that is similar to PVF coverage defects. This work establishes a requirement for PDGFRα signaling in PVF development and may shed light upon the potential pathways that are over-activated in PVFs in injury and disease contexts.

## Introduction

The anterior forebrain is vascularized early in mouse development (embryonic day or E10) by a blood vascular network that first appears in progenitor domains like the ventricular zone (Vasudevan et al., 2008). Soon after, at ∼E12, penetrating vessels from the perineural vascular plexus in the meninges grow inward to vascularize superficial regions like the neocortical plate that eventually is part of the cerebral cortex as the neocortex (Stubbs et al., 2009). The spatiotemporal process of anterior forebrain vascularization is similar in larger mammals, including rabbits (Strong, 1964) and humans (Marín-Padilla, 2012). This early stage of cerebrovascular development is the initial steps in a process that spans pre- and postnatal development to generate the dense network of hierarchically-organized cerebral vessels. The cerebrovasculature enables constant delivery of oxygen and nutrients via the blood while protecting the cerebral parenchyma, via the blood-brain barrier (BBB), from disruptive peripheral blood contents. Disruptions to pre- and postnatal stages of cerebrovascular development can negatively impact brain development (reviewed in Ouellette et al., 2024; Ouellette and Lacoste, 2021; Wälchli et al., 2023) and, later in life, diminished cerebrovascular function and integrity is a hallmark feature of age-related neurodegenerative disorders (reviewed in Sweeney et al., 2018). Therefore, comprehensive knowledge of the mechanisms underlying cerebrovasculature formation and maturation has important implications for brain disorders across the lifespan.

A key part of building the cerebrovascular network is the maturation of the cell types that comprise the neurovascular unit (NVU). The NVU is made up of specialized cell types that coordinate to form a fully functional vascular system, including connected endothelial cells that make up the blood vessel lumen and perivascular cells including mural cells (vascular smooth muscle cells or vSMC and pericytes), the endfeet processes of astrocytes, perivascular macrophages (PVMs) and perivascular fibroblasts (PVFs). NVU perivascular cells have separate but overlapping maturation timelines. In mice, mural cells that directly contact endothelial cells co-develop with the prenatal vasculature (Daneman et al., 2010; Jung et al., 2018). During perinatal-postnatal development, mural cells upregulate markers of mature vSMC (contractile proteins alpha-smooth muscle actin and Myh11) that surround large diameter arterioles and mature pericytes (Anpep/CD13) on small diameter capillaries (Coelho-Santos and Shih, 2020; Slaoui et al., 2023). In contrast to mural cells, astrocytes are not born until late stages of mouse cerebral development and therefore ‘join’ the NVU later than mural cells. Astrocyte endfeet are contacting vessels at birth in mice but substantially refine their contact, morphology and polarization of endfeet proteins like Aquaporin-4 (AQP-4) during the first two postnatal weeks (Freitas-Andrade et al., 2023; Munk et al., 2019). PVMs and PVFs differ from mural cells and astrocytes in that they are only localized around large diameter, non-capillary cerebral vessels. PVMs and PVFs are the last NVU cells to appear developmentally on the cerebrovasculature. They migrate from the meninges into perivascular spaces around penetrating arterioles and (to a lesser extent) ascending venules, expanding to cover vessels to their termini via proliferation and migration between P5 and P14 (Jones et al., 2023; Karam et al., 2022; Masuda et al., 2022). PVM and PVF cell bodies are between the vSMC layer and astrocyte endfeet. We previously showed PVMs and PVFs develop in tandem (Jones et al., 2023), suggesting potential cross-talk between these cells during development. To varying degrees, we know the molecular mechanisms and signaling pathways underlying development of endothelial cells, mural cells, astrocyte and PVM (Daneman et al., 2009; Freitas-Andrade et al., 2023; Hellström et al., 1999; Hogan et al., 2004; Liebner et al., 2008; Masuda et al., 2022; Munk et al., 2019; Stenman et al., 2008; Zhou and Nathans, 2014), however we currently lack this knowledge for PVFs.

PVFs are distinguishable from other cell types in the NVU by expression of collagen 1 and expression of both platelet-derived growth factor receptors (PDGFRs), PDGFRα and PDGFRβ, characteristics they share with fibroblasts in the meninges (Dorrier et al., 2021; Kelly et al., 2016; Pietilä et al., 2023; Soderblom et al., 2013; Vanlandewijck et al., 2018). PDGFRs are receptor tyrosine kinases that activate by formation of homodimers or heterodimers upon binding of PDGF ligands AA, BB, CC, or DD. Within the CNS, PDGFR signaling plays important roles in glial and vascular development, differentiation, and maintenance (Sil et al., 2018). PDGFRα is expressed by oligodendrocyte precursor cells (OPCs), and PDGFRα/PDGF-AA signaling is involved in OPC proliferation, survival, and differentiation (Cardona et al., 2021; Pringle et al., 1989; Richardson et al., 1988). PDGF-BB ligand is produced by the endothelium and is required for the recruitment and maintenance of PDGFRβ-expressing mural cells during CNS vascular development (Hellström et al., 1999; Lindahl et al., 1997; Lindblom et al., 2003). *Pdgfc*-null mice that are additionally heterozygous for a *Pdgfra*-null allele have severe defects in meningeal basement membrane deposition and cortical development, suggesting a role for PDGFRα/PDGF-CC in the proper development and function of CNS fibroblasts (Andrae et al., 2016). Collectively, these studies raise the possibility that PDGF signaling plays a role in PVF development.

Here we provide evidence from *in vivo* approaches that PDGFRα is required in PVFs for proper coverage of vessels during postnatal development. Most perivascular NVU cell types and structures are not overtly altered by perturbation in PVF development. The exception is PVMs, which show a similar reduction in coverage as PVFs, pointing to an important role of PVFs in supporting development of PVMs in the cerebrovasculature.

## Results

### PVFs express PDGFRα and PDGFRβ during postnatal development

CNS fibroblasts, including PVFs and meningeal fibroblasts, express both PDGFR proteins (Kelly et al., 2016; Soderblom et al., 2013). We sought to validate these findings in PVFs at both the protein and RNA levels. Immunostaining shows expression of PDGFRα and PDGFRβ within collagen-1-positive cells in the meninges and cortex of P21 wild-type mice (Fig. 1A, arrowheads). Expression of these receptors is also observed in the expected cell types as defined by location and morphology: OPCs are enriched for expression of PDGFRα (Fig. 1A, arrows), and pericytes show expression of PDGFRβ (Fig. 1B, arrows). We next examined PDGFR expression at the transcript level. Kelly et al., 2016 showed protein expression of PDGFRα and β within CNS fibroblasts/PVFs at P0 and P21, our work shows PVFs emerge on vessels between P5 and P14 (Jones et al., 2023). Thus, we looked at expression of both PDGFRs at P8, in the middle of the PVF developmental window, using the *Collagen1a1(Col1a1)-GFP* transgenic mouse line which is an established reporter line for labeling CNS fibroblasts (Bonney et al., 2022b; Jones et al., 2023; Soderblom et al., 2013). Using RNAScope *in situ* detection, we observe expression of both *Pdgfra* and *Pdgfrb* transcript within *Col1a1-GFP+* meningeal fibroblasts and PVFs on vessels in the cortex, with additional expression in the expected cell types (*Pdgfra* in OPCs, *Pdgfrb* in vSMCs and pericytes) (Fig. 1C). These experiments establish dual expression of PDGFRs by PVFs during the period they develop and cover the parenchymal vasculature.

**Figure 1:**
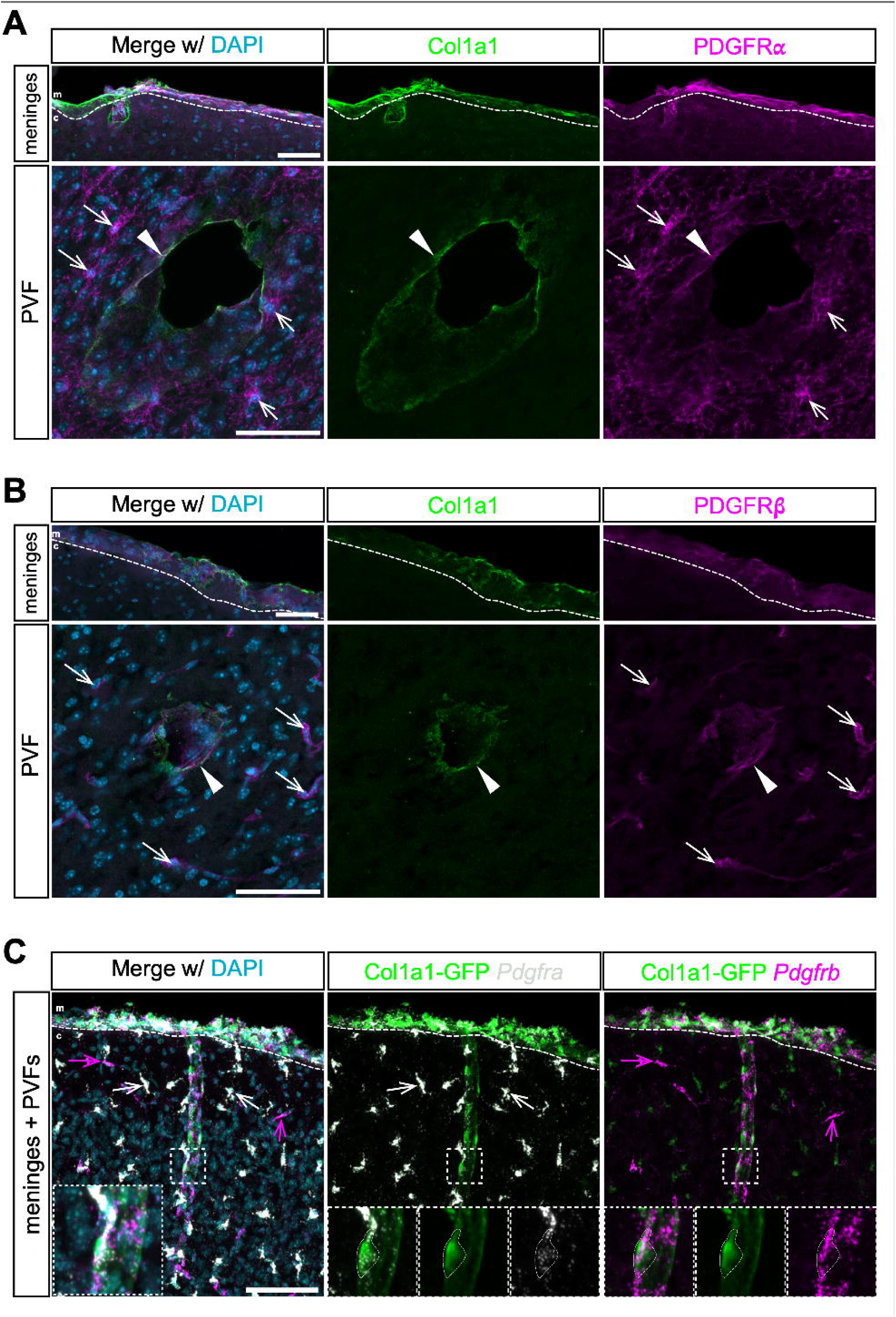
PVFs express PDGFRs during their development. (**A-B**) Confocal images of meninges and cortical PVFs in P21 wildtype-control (Col1a2-Cre^+/+^) mice, showing protein staining for Col1a1 (green) and (**A**) PDGFRα (magenta) and (**B**) PDGFRβ (magenta). **m**: meninges, **c**: cortex; dashed line denotes border between meninges and cortex. Arrows denote (**A**) OPCs expressing PDGFRα and (**B**) pericytes expressing PDGFRβ; arrowheads denote PVFs. Scale bars 50μm. (**C**) Confocal image of meninges and penetrating vessel in P8 *Col1a1-GFP* mouse, showing RNAscope transcript detection of *Pdgfra* (white) and *Pdgfrb* (magenta) in *Col1a1-GFP+* PVFs and meningeal fibroblasts (green). **m**: meninges, **c**: cortex; dashed line denotes border between meninges and cortex; dashed boxes indicate area of inset. Arrows denote OPCs expressing *Pdgfra* (white arrows) and pericytes/vSMCs expressing *Pdgfrb* (magenta arrows). Scale bar 100μm.

### Expression patterns of PDGF ligands in the postnatal brain

PDGFR dimers display specificity for certain ligand combinations, based on phenotypic data from knockout models of the receptors or ligands as well as *in vitro* findings in a range of mesenchymal cell types. PDGFRα homodimers are bound and activated by PDGF-AA and PDGF-CC, while PDGFRα/β heterodimers and PDGFRβ/β homodimers are primarily bound and activated by PDGF-BB (Fig. 2A) (Boström et al., 1996; Campaña et al., 2025; Ding et al., 2004; Fantauzzo and Soriano, 2017, 2016; Levéen et al., 1994; Soriano, 1997, 1994).Widespread expression of PDGF ligands has been reported within the developing mouse forebrain. PDGF-A and -C are expressed in the dorsal cortex beginning around birth (Andrae et al., 2016), PDGF-B is present at high levels in the endothelium embryonically through birth (Hellström et al., 1999) and lower, sustained expression postnatally through adulthood (Vazquez-Liebanas et al., 2022). PDGF-D ligand expression in brain development is not widely reported, except in the thalamus and floor plate of the pontine area in embryonic rat brains (Hamada et al., 2002). Since PVFs express both PDGFRα and PDGFRβ during their postnatal development, we tested which ligands are present that might act on PVFs, focusing on *Pdgfa*, *Pdgfb*, and *Pdgfc* in *Col1a1- GFP+* P8 brains. We see expression of *Pdgfa* and *Pdgfc* within the brain within what are likely neurons. There is some expression of *Pdgfb* and *Pdgfc* observed in the meninges, likely attributed to expression by meningeal blood vessels and vSMCs, respectively based on single cell data sets of vascular cells (Vanlandewijck et al., 2018) (Fig. 2B, D). We observe enrichment of *Pdgfb* expression specifically within vessels, including large penetrating vessels containing PVFs (Fig. 2C, outlines). These results confirm expression of PDGF-AA, -BB and -CC ligands within the temporal window of PVF migration, though only PDGF-BB appears to be enriched in the vasculature near developing PVFs.

**Figure 2:**
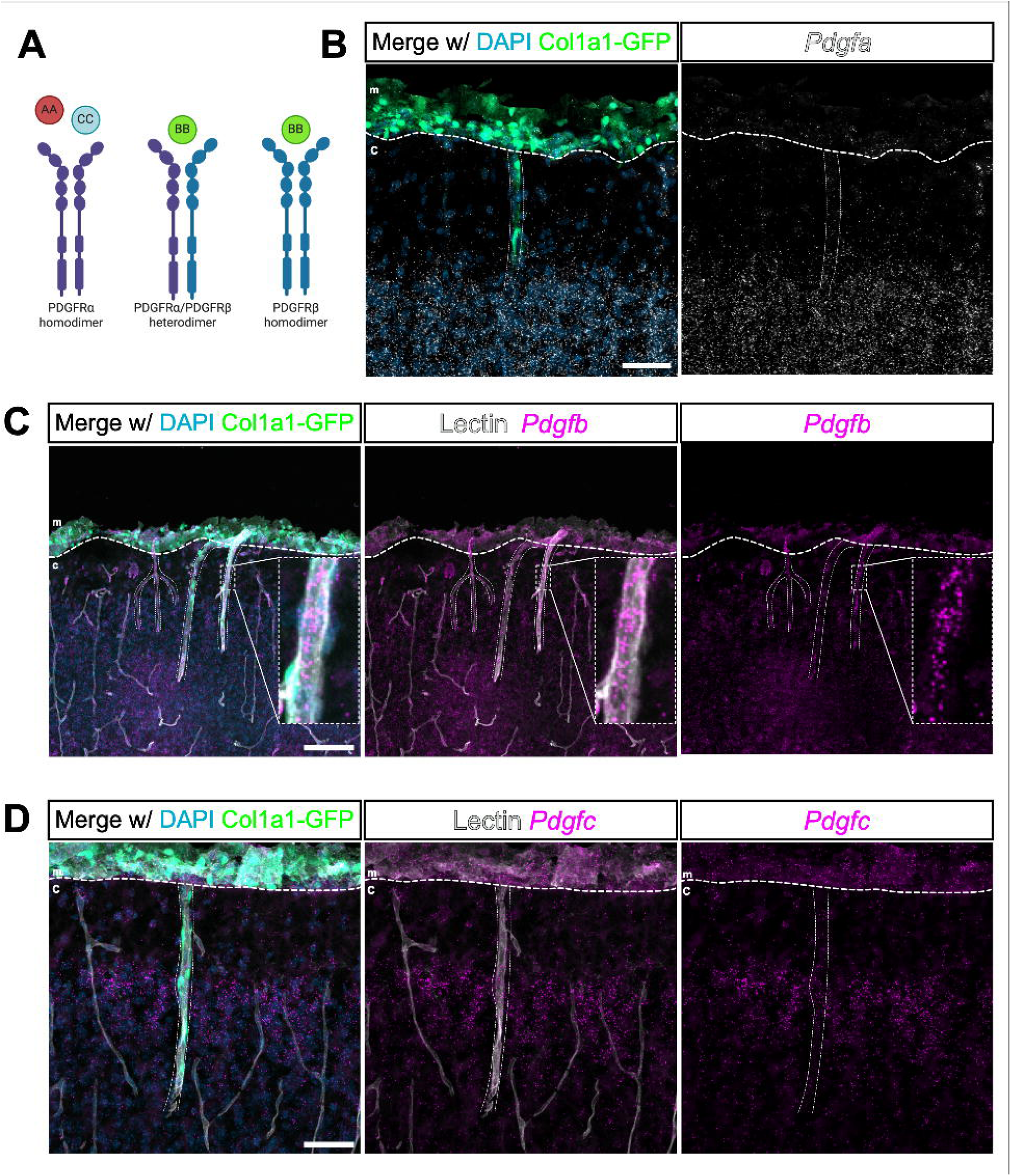
Expression patterns of PDGF ligands A, B, and C in the postnatal brain. (**A**) Diagram illustrating PDGF ligand-receptor pairs. (**B**-**D**) Confocal images of meninges and penetrating vessels in a P8 *Col1a1-GFP* mouse, showing *Col1a1-GFP+* PVFs and meningeal fibroblasts (green) and RNAscope transcript detection of (**B**) *Pdgfa* (white), (**C**) *Pdgfb* (magenta), and (**D**) *Pdgfc* (magenta); nuclei labeled with DAPI (blue). Fine dashed lines outline vessels, in (**C**-**D**) labeled with lectin (white). **m**: meninges, **c**: cortex; larger dashed lines denote border between meninges and cortex. Scale bars (**B,D**) 50μm, (**C**) 100μm.

### Reduction in PDGFRα expression by fibroblasts perturbs PVF coverage and distribution on cerebral vessels

Given the overlapping expression of PDGFRβ in mural cells and PVFs, we opted to delete PDGFRα just prior to postnatal PVF development, as this was the only PDGFR expressed exclusively in this perivascular cell type. To do this, we employed a tamoxifen-inducible genetic model to conditionally delete *Pdgfra* in fibroblasts. Specifically, we first crossed *Pdgfra-H2B- GFP* mice, in which a histone-targeted GFP inserted into the endogenous locus results in a null allele, with *Col1a2-CreERT* mice, which drive Cre expression in CNS fibroblasts (Bonney et al., 2022b; Dorrier et al., 2021; Jones et al., 2023), to generate *Col1a2-CreERT^Cre/+^; Pdgfra-H2B-GFP^GFP/+^* mice. Notably, *Col1a2-CreERT* mice have very minimal recombination frequency in vSMCs (<1%), however, this cell type does not express PDGFRα and is easily distinguishable from fibroblasts based on its distinct morphology (Bonney et al., 2022b; Vanlandewijck et al., 2018). We adopted this approach after observing that conditional deletion of *Pdgfra* using only floxed alleles did not result in efficient recombination and protein reduction when crossed with *Col1a2-CreERT* (data not shown), highlighting the necessity of introducing the *H2B-GFP* null allele for effective recombination. We generated conditional knockout (cKO) mutant mice by crossing *Col1a2-CreERT^Cre/+^; Pdgfr*α*-H2B-GFP^GFP/+^* mice with *Pdgfra-flox^fl/fl^* mice also expressing a *Ai14(tdTomato)-flox* reporter allele, allowing for visualization of tdTomato+ Cre recombined cells upon tamoxifen recombination (Fig. 3A). To examine how partial or complete *Pdgfra* loss affects PVF development and vascular coverage in the cortex, we compared cKO mice to three additional genotypes: (1) control for the *Pdgfra* allele (*Col1a2-CreERT^Cre/+^; Ai14^fl/+^; Pdgfra^+/+^*), (2) global heterozygous (gHet) mice (*Col1a2-CreERT^Cre/+^; Ai14^fl/+^; Pdgfra- H2B-GFP^GFP/+^*), which retain one functional *Pdgfra* allele and allow for visualization of tdTomato+ recombined fibroblasts and GFP+ PDGFRα-expressing cells, and (3) conditional heterozygous (cHet) mice (*Col1a2-CreERT2^Cre/+^; Ai14^fl/+^; Pdgfra-flox^fl/+^*), which have a single allele of PDGFRα recombined in Cre-expressing tdTomato+ CNS fibroblasts upon tamoxifen exposure (Figure 3A).

**Figure 3.**
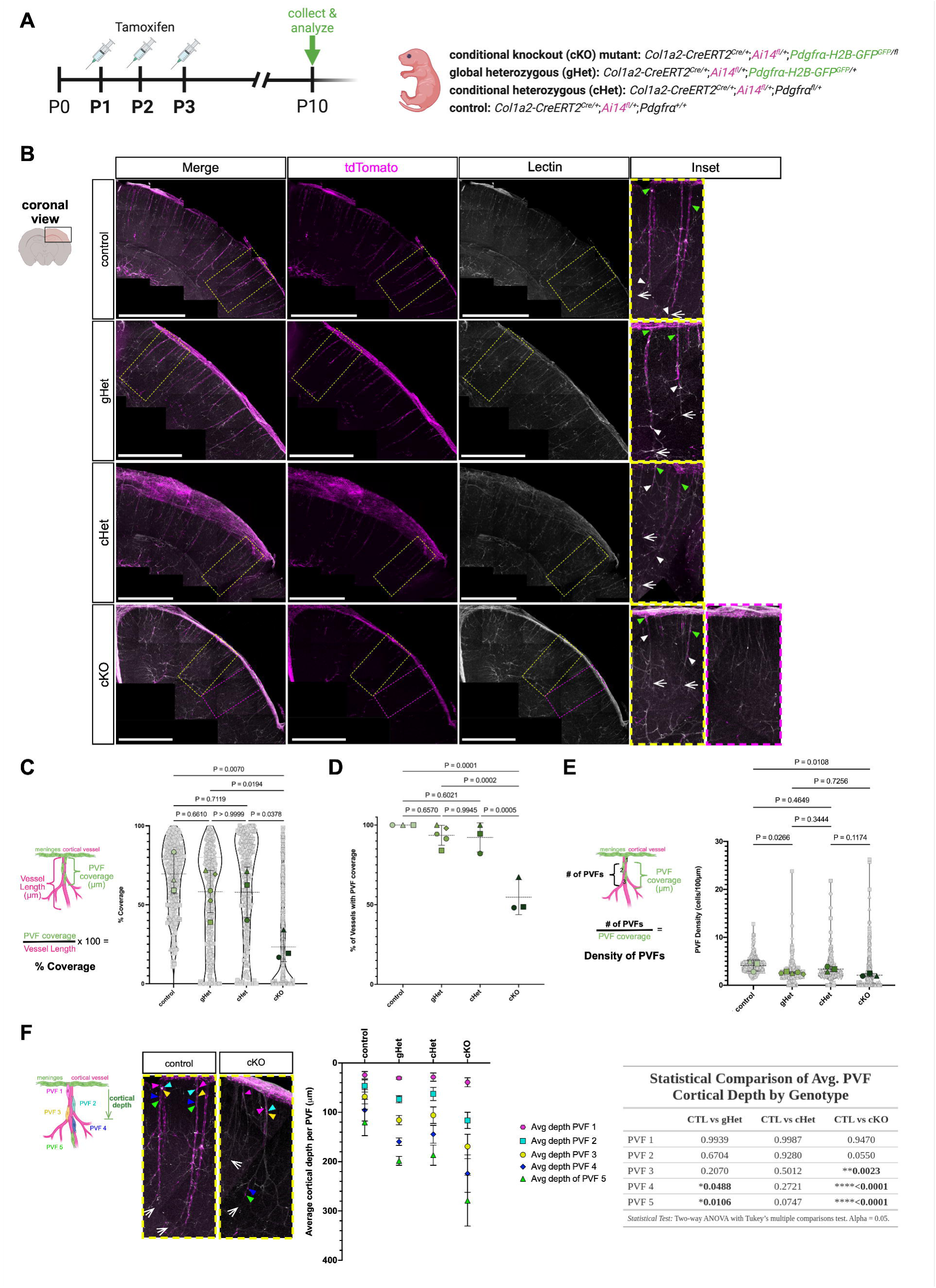
Reduced PDGFRα signaling impairs PVF coverage of penetrating cerebral vessels blood vessels during postnatal development. (**A**) Schematic of experimental design including timeline for tamoxifen injections and P10 collection. (**B**) Representative maximum projection images of coronal sections (z-volume: 500 – 1000µm) of P10 CUBIC-cleared brains carrying the tamoxifen-inducible Cre reporter allele *Col1a2-creERT2^Cre/+^; Ai14(tdTomato)- Flox^fl/+^* showing fibroblasts (RFP, magenta) and vessels (Lectin, white). White arrows in insets mark vessel terminus. Green carets mark emerging PVFs. White carets indicate the cortical depth of the furthest PVF on a blood vessel. Scale bars: 1 mm. (**C**) Graph depicts percent coverage of vessels by PVFs at P10. Data are mean±s.d. The mean for each biological replicate (*n*=3-5 for each genotype) is represented by large colored shapes and the corresponding technical replicates (vessels per animal) in small gray matching shapes. One-way ANOVA with multiple comparisons indicates statistically significant differences in percent coverage between: control vs. cKO, *P=0.0070*; gHet vs. cKO, *P=0.0194*; cHet vs. cKO, *P=0.0378*. (**D**) Graph depicts percent of penetrating cerebral vessels with any PVF coverage at P10. One-way ANOVA with multiple comparisons indicates statistically significant differences in percent coverage between: control vs. cKO, *P=0.0001*; gHet vs. cKO, *P=0.0002*; cHet vs. cKO, *P=0.0005*. (**E**) Graph depicting PVF density (PVFs per 100um vessel length) on large diameter cortical penetrating vessels at P10. Data are mean±s.d. The mean for each biological replicate (*n*=3-5 for each genotype) is represented by large colored shapes and the corresponding technical replicates (vessels per animal) in small gray matching shapes. One-way ANOVA with multiple comparisons indicates statistically significant differences in percent coverage between: control vs. cKO, *P=0.0108*; control v.s. gHet, *P=0.0266*. (**F**) Representative images of control and cKO vessels in which PVFs 1-5 are depicted by colored arrows. Line in cKO indicates a gap in PVF coverage. Schematic of analysis and graph depicting average cortical depth of ranked PVFs 1-5 per genotype. Data are mean±s.d. Table reports statistical tests and p-values, reporting comparison to control (CTL).

We assessed recombination efficacy and PDGFRα expression in P10 mice following tamoxifen administration at P1-3 (Fig. 3A, Sup. Fig. 1). Immunostaining reveals a qualitative decrease in PDGFRα expression in the meninges and along penetrating cortical vessels in gHets compared to controls and no detectable protein expression in cKO mutants (Sup. Fig. 1A). Quantification of the number of tdTomato+ meningeal fibroblasts per 100μm length of meninges shows no differences in meningeal density across all genotypes (Sup. Fig. 1B). Fluorescent RNA *in situ* hybridization shows a qualitative reduction in *Pdgfra* transcript levels in cKO mutants compared to controls (Sup. Fig. 1C). We also examined other PDGF pathway components by performing RNA *in situ* hybridization for *Pdgfrb* and *Pdgfb* in control, cHet, gHet, and cKO mice at P10. *Pdgfrb* expression by PVFs and *Pdgfb* expression by the endothelium is unchanged across genotypes (Sup. Fig. 1D-E), consistent with previous data that loss of PDGFRα does not affect expression of PDGFRβ (Fantauzzo and Soriano, 2016). This analysis demonstrates effective postnatal deletion of PDGFRα in cKO without overt impact on expression of other relevant PDGF ligand and receptor components in the NVU.

We assessed PVF coverage and depth on vessels in the four genotypes at P10. We selected P10 based on our prior work establishing a time course of PVF development between P5 and P14. At P10, PVFs have ∼75% coverage of the cerebral vessel lengths but are still actively developing as PVFs reach near complete coverage by P14 (Jones et al., 2023). We performed CUBIC tissue clearing on 2 mm brain slices, followed by labeling with tomato-lectin to visualize the vasculature. In controls, gHets and cHets, PVFs are distributed along penetrating cerebral vessels from their meningeal origin at the pial surface to their termini in deeper cortical regions (Fig. 3B). In controls, PVFs are near the vessel termini (Fig. 3B, insets) however in the gHets and cHets, we observe some PVFs farther from the vessel termini (Fig. 3B, insets). In cKO mutants, we observe regions with partial coverage and regions in which no cerebral vessels had PVFs (Fig. 3B, yellow and magenta boxes denote magnified insets). Insets from areas of the cKO with partial coverage show the last PVFs on the vessel are situated far from the vessel termini (Fig. 3B, yellow insets).

We quantified percent PVF coverage of individual vessels in the four genotypes, determined by measuring the length of the PVF-labeled region relative to total vessel length (Fig. 4C). Control mice exhibit the highest average PVF coverage (69.15%), whereas cKO mutants show a significant reduction (23.27%; p = 0.0070). While gHets and cHets have PVF coverage similar to control, cKO mutants have significantly less coverage than both gHets (p = 0.0194) and cHets (p = 0.0378). We also quantified the percent of penetrating cerebral vessels in each genotype with any PVF coverage, which reveals cKO have significantly fewer cerebral vessels with any PVFs as compared to control, gHet and cHet (Fig. 3D).

**Figure 4:**
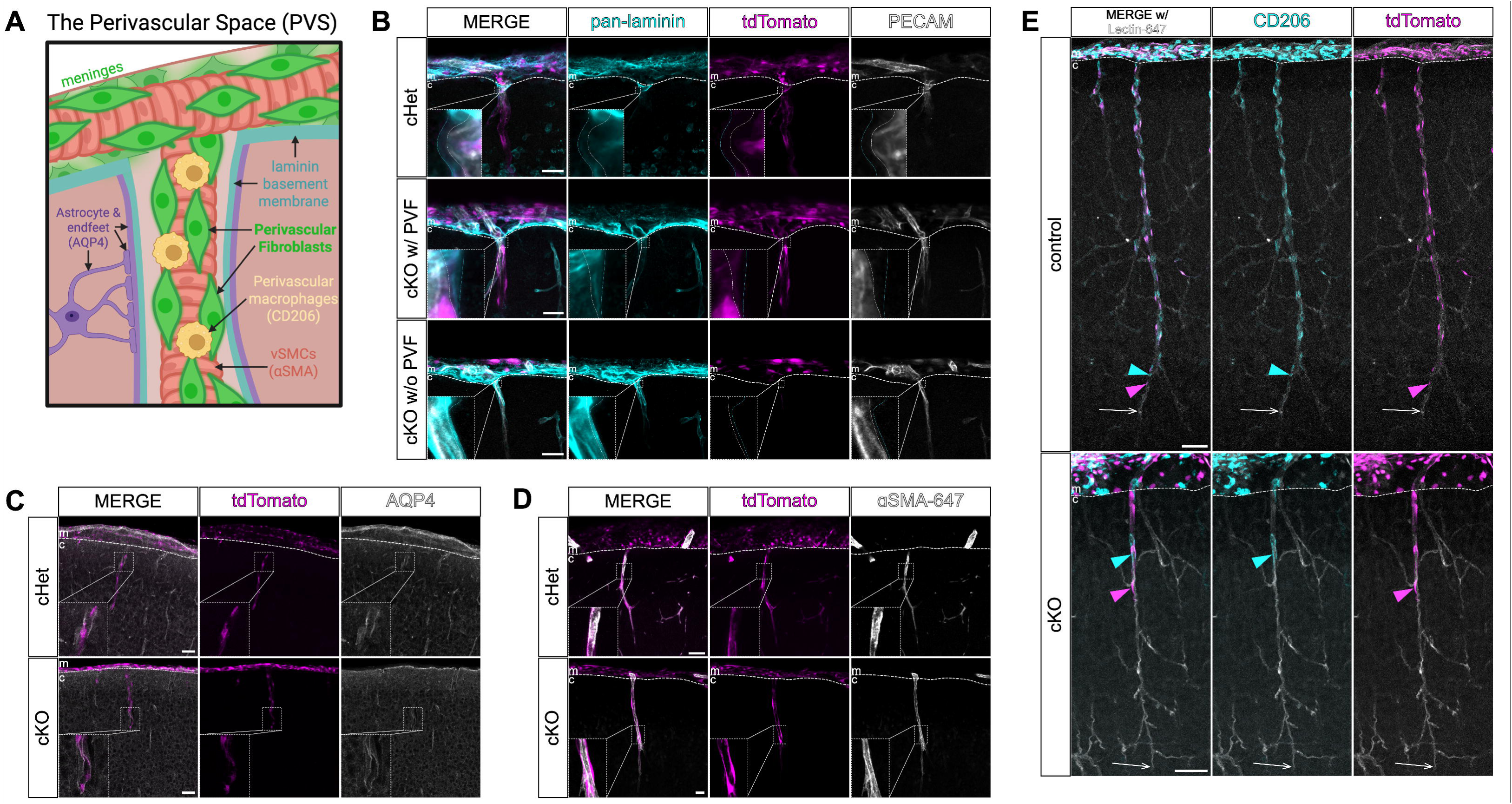
PDGFRα cKO from PVFs alters PVM development but not other features of the perivascular space. (**A**) Graphical depiction of the perivascular space (PVS) showing blood vessels ensheathed by vSMCs (pink), astrocytes and their endfeet (purple), basement membrane (cyan), perivascular macrophages (yellow) and perivascular fibroblasts (green). (**B**-**E**) Confocal images of meninges and penetrating vessels in control (cHet or control) and cKO mutant mice at P10 showing fibroblasts labeled with tdTomato (magenta), with protein staining for PVS structures and cell types: (**B**) basement membrane labeled by pan-laminin (cyan) and blood vessels labeled by PECAM (white); (**C**) astrocytes and endfeet labeled by AQP4 (white); (**D**) vSMCs labeled by αSMA (white); and (**E**) macrophages labeled by CD206 (cyan) and blood vessels labeled by lectin (white). **m**: meninges, **c**: cortex; dashed line denotes border between meninges and cortex. Areas of inset indicated by dashed boxes. Dashed lines in (**B**) insets outline basement membrane (cyan) and vessel surface (white). White arrows in (**E**) mark vessel terminus, magenta and cyan carets indicate the cortical depth of the furthest migrated PVF and PVM, respectively, on vessel. Scale bars (**B**, **D**) 25μm, (**C**, **E**) 50μm.

We also looked at PVF density and distribution at P10 in cerebral vessels with PVF coverage. We analyzed PVF density (PVF # per 100μm) within covered regions. gHets and particularly cKO mutants exhibit a significant reduction in PVF density as compared to control (Fig. 3E). Upon qualitative assessment of control and cKO penetrating cerebral vessels, we noticed PVFs in controls cover vessels at even, regularly-spaced intervals between PVF cell bodies. This contrasted with often large, PVF-free gaps between individual PVFs in cKOs (Fig. 3F, images). To systematically evaluate these gaps across genotypes, we measured the average depth of the cell body of the first five PVFs along individual vessels (Fig. 3F). This approach was based on the observation that cKO mutant vessels with coverage had at least five PVFs, permitting sufficient vessel numbers for analysis. Our analysis reveals a significant increase in the average depth of PVFs 3-5 in cKO mutants compared to control and PVFs and PVFs 4 and 5 in gHets (Fig. 3F, graph and table).

These findings support that cKO PVFs that completely lack PDGFRα during PVF development are less likely to enter the perivascular space and if PVFs enter, they do not sufficiently expand in number, leading to reduced vessel coverage. Consistent with this, partial or complete absence of PDGFRα results in altered distribution and decreased density of PVFs on vessels. Collectively this data points to an important role for PDGFRα in PVF proliferation, survival and/or migration during PVF development.

### Impact of decreased fibroblast PGDFRα expression on the perivascular niche

We next tested the impact of reduced PVF coverage on other perivascular cell types in the NVU we conducted immunostaining to look at structural components and cell types of the NVU, including basement membrane, astrocytic endfeet, vSMCs, and PVMs (Fig. 4A). Laminins are a critical component of the vascular, pial and parenchymal basement membranes (Hannocks et al., 2018; Nirwane and Yao, 2022). To test if the basement membrane is affected in cKO mutants, we immuostained cHet (no difference in PVF coverage from control) and cKO mutant brains at P10 for pan-laminin and PECAM to visualize basement membrane and the vessel wall, respectively (Fig. 4B). In cHet and cKO we observe continuous pan-laminin labeling of the basement membrane at the pial surface and the vascular basement membrane co-localized by PECAM on vessels with and without PVFs (Fig. 4B). Insets highlight where the vessel enters the parenchyma to show a small gap between the parenchymal and vascular basement membranes in cHet and cKO with and without PVF (Fig. 4B, insets). Prior work using laminin and collagen immunolabeling and electron microscopy show the PVS is wider and more visible near the vessel entry point into the brain and a short distance past the pial surface (Bonney et al., 2022; Hannocks et al., 2018; Masuda et al., 2022). Collectively, this indicates the vascular, pial and parenchymal basement membranes overtly intact in cKO mutants, including at entry points of vessels into the brain at this stage of development.

Another critical developmental event required to form the PVS is the maturation of astrocytic endfeet, which occurs postnatally with the polarization of AQP4 to endfeet (Freitas-Andrade et al., 2023; Munk et al., 2019). We assessed if astrocytic endfoot maturation is overtly disrupted in cKO mutants by conducting immunostaining for AQP4 in cHet and cKO mutant brains at P10. We see no differences in AQP4 localization or coverage on vessels between cHet and cKO mutants (Fig. 4C). This suggests astrocyte endfoot development and maturation is not obviously impacted by reduction or lack of PVF coverage. vSMCs wrap large penetrating arterioles in the brain and are responsible for regulating vascular blood flow. vSMCs are established around the vasculature prior to the development of PVFs (Slaoui et al., 2023), thus we sought to understand if vSMC coverage is altered in cKO mutants and may underly lack of PVF coverage. We conducted immunostaining for vSMCs in control (cHet) and cKO mutant brains at P10 using a directly-conjugated secondary antibody against αSMA and see no differences in vSMC coverage on control versus mutant vessels (Fig. 4D). On cKO vessels that lacked PVF coverage, we still observe vSMCs (Sup. Fig. 3A). Collectively, this data suggests the impairment in PVF coverage seen in cKO mutants is due to cell-intrinsic loss of PDGFRα leading to failure of PVFs to proliferate, survive and/or migrate within PVSs. Likewise, loss or reduced numbers of PVFs on cKO vessels does not appear to overtly affect PVS components such as basement membrane, astrocyte endfeet, or vSMC coverage at this stage of development.

Previous work from our group and others showed PVMs cover parenchymal vessels postnatally and we show they develop concurrently with PVFs (Jones et al., 2023; Karam et al., 2022; Masuda et al., 2022). Given the severity of the PVF coverage defect in cKO mutants we next tested if PVM coverage is equally impaired, or if, conversely, PVM development can proceed normally independent of PVFs. To test this, we immunostained control and cKO mutant CUBIC-cleared 2 mm brain slices at P10 with CD206 to label PVMs and tomato-lectin to visualize the vasculature. On control vessels, PVM coverage matched the extent of PVF coverage, with both cell types evenly covering the length of vessels from the pial surface nearly to the vessel terminus (Fig. 4E). On cKO vessels PVM coverage was overtly reduced and matched the reduced pattern of PVF coverage, with large stretches of vessel uncovered by both PVFs and PVMs (Fig. 4E, Sup. Fig. 2B). This data, together with our prior work, suggests a requirement for PVFs in guiding PVM development.

## Discussion

Our work identifies PDGF signaling pathway components are expressed during PVF development and PDGFRα expression by PVFs as required for initial vascular coverage in the postnatal cerebral cortex. We also show that altered PVF coverage of vessels impairs initial cerebrovascular coverage of CD206+ PVMs in the NVU. Collectively, our observations provide important new insights into the signaling pathways that control NVU development and previously unappreciated cross-talk between NVU cell types, fibroblasts and macrophages.

PDGF ligand and receptors have known functions in CNS development, and specifically in assembly of the NVU (Sil et al., 2018). PDGFRβ/PDGF-BB signaling is required for mural cell recruitment to the endothelium during CNS vascular development (Hellström et al., 1999; Lindahl et al., 1997; Park et al., 2017) and maintenance of mural cells in the adult CNS (Vazquez-Liebanas et al., 2022). Consistent with other reports using a *Pdgfb* Cre reporter (Claxton et al., 2008), we also observe *Pdgfb* expression by the endothelium at P8 during the window of PVF development. Work using the PDGF-BB *ret* mutant, in which the retention motif that tethers PDGF-BB to the basement membrane of vessels is deleted, shows that tethering of PDGF-BB to the vasculature is required for mural cell recruitment (Lindblom et al., 2003). Possibly, endothelial PDGF-BB acts as a ‘local’ signal to stimulate expansion on cerebral vessels. PDGF-A and PDGF-C expression was observed more broadly in the brain, likely being expressed by neurons, and some low-level expression in the meninges, potentially by vSMC that based on scRNAseq data sets express *Pdgfa* and *Pdgfc* (Vanlandewijck et al., 2018). Therefore, PDGF-AA and PDGF-CC ligands from multiple sources could activate the PDGFRα receptor on PVFs to stimulate proliferation, migration and/or survival. In support of this, mice with combined mutations in *Pdgfc* and *Pdgfra* have defective meningeal fibroblast development (Andrae et al., 2016) and we have shown previously that PVFs arise from fibroblasts in the meninges. Future experiments using mouse mutants as well as *in vitro* approaches are required to determine the sufficiency of PDGF ligands to drive PVF development. Additionally, it is currently unknown which PDGFR dimers form and are active in PVFs. Future experiments using bimolecular fluorescence complementation (Campaña et al., 2025; Rogers et al., 2022) could be used to determine the specific PDGFR dimer pairs that form in PVFs, and further point towards the specific ligand-receptor pairs and downstream signaling pathways involved in PVF development.

Conditional deletion of PDGFRα from CNS fibroblasts postnatally leads to fewer cerebral vessels with any PVF coverage and reduced PVF coverage of vessels at P10. This points to an impairment in the initiation of PVF coverage of cerebral vessels (P5-P7) and expansion of PVF number on cerebral vessels via local proliferation, survival and/or migration (P7 and P14). PVFs are detected on leptomeningeal vessels, which are connected to the cerebrovasculture, in embryonic development yet they do not appear on cerebral vessels until P5-P7. What triggers this initiation of coverage of parenchymal vessels is not known, however our data support that PDGFRα-mediated signaling is a critical part of this process. We did not observe overt changes in other perivascular NVU cell types that are established prior to the appearance of PVFs and thus could influence PVF development, such as astrocyte endfeet polarization and vSMC coverage. Prior work on PVM development identified perivascular gaps between the vessel wall and the laminin-enriched glial limitans where leptomeningeal vessels enter the brain parenchyma between P4 and P9 (Masuda et al., 2022). The appearance of these gaps coincides with both PVF and PVM appearance around cerebral vessels (Jones et al., 2023; Masuda et al., 2022). This indicates that access to penetrating vessels is one part of the initiation of PVF and PVM coverage of cerebral vessels. We observed perivascular spaces where leptomeningeal vessels enter the brain parenchyma even on vessels that do not contain PVFs in fibroblast cKO of PDGFRα, suggesting that access to cerebral vessels is not overtly altered. This supports the primary defect underlying impaired initiation of PVF coverage of cerebral vessels is loss of PDGFRα-mediated signaling in PVFs. A plausible model is that PVFs that lack PDGFRα have reduced response to one or more PDGF ligands, thus decreasing PVF survival and/or migration, and are therefore less likely to initiate coverage of cerebral vessels.

We previously showed using clonal analysis that after initiation of coverage, single PVFs locally proliferate and migrate to cover portions of a single vessel (Jones et al., 2023). Our observation of reduced cerebrovascular coverage and PVF density in PDGFRα cKO mutants supports that, for PVFs that do initiate coverage, local proliferation and/or survival on vessels is likely impaired. This is also supported by our distribution analysis of the first five PVFs on individual vessels that showed PVFs 3-5 were deeper on cerebral vessels in global hets and cKO. This suggest that PVFs in cKO mutants retain some capacity to migrate on the vasculature but their expansion in number via proliferation and/or survival is impaired, leading to PVF-free gaps. The stepwise increase in depth at PVF positions 3-5 may indicate a compensatory attempt by existing PVFs to migrate deeper along vessels in the absence of sufficient proliferation. The failure to establish continuous PVF coverage supports that PDGFRl7l signaling is required not only for the initial positioning of PVFs along vessels but also for their expansion via proliferation and/or survival. These potential roles for PDGFRα signaling in the context of PVFs will be explored in future experiments.

The NVU contains PVMs that we showed previously develop in tandem with PVFs (Jones et al., 2023). In PDGFRα cKO mutants with reduced PVF coverage at P10, PVM distribution is qualitatively altered with macrophages not extending past the deepest PVF and showing similar gapped coverage as PVFs. This indicates that PVMs depend on PVFs for their initial development on cerebral vessels. Recent work showed that IL-34, a ligand for colony-stimulating factor 1 receptor (CSF1R) expressed by macrophages, is required for perivascular macrophage coverage of vessels and that IL-34 produced by mural cells and fibroblasts in the NVU is a major source (Hove et al., 2025). Deletion of talin, a cytoskeletal protein required for integrin mediated cell adhesion, migration and signaling (Calderwood et al., 2013), from macrophages impairs PVM density (Masuda et al., 2022). Integrin-mediated migration occurs via binding to ECM components such as fibronectin, collagen and vitronectin, which are expressed to varying degrees by endothelial cells, mural cells and PVFs (Hannocks et al., 2018; Pietilä et al., 2023; Vanlandewijck et al., 2018). This data suggests that PVFs, potentially through production of IL-34, a mitogen and pro-survival signal for macrophages via activation of CSF1R, and deposition of ECM components like collagen, is a key NVU cell type for initial PVM coverage of vessels. More studies looking at later timepoints are required to test if PVM development is only delayed in PDGFRα cKO or if PVFs play a role in PVM maintenance in the adult CNS.

While the focus of this study is PVF development on cerebral vessels, our identification of a key role for PDGFRα aligns with work implicating PDGF signaling in CNS fibroblast response to brain injury and disease. PVFs are implicated in fibrosis in CNS disease and injury (Dorrier et al., 2022; Fernández-Klett et al., 2013; Kelly et al., 2016; Månberg et al., 2021; Soderblom et al., 2013). These cells thus represent a potential therapeutic target for modulating the severity of disease and injury sequelae. Studies targeting PDGF ligand/receptor interactions following CNS injury have been shown to reduce injury severity, promote axon regeneration, and restore blood-brain barrier properties (Nguyen et al., 2021; Yao et al., 2022). Inhibition of PDGFRα was recently shown to inhibit expansion of pro-fibrotic myofibroblasts following cerebral ischemia and improve stroke recovery (Protzmann et al., 2025). This sets up the intriguing possibility that PDGFRα plays dual roles in PVF development and is part of the fibrotic response following CNS injury. Understanding the role of PDGFRs in PVFs during development and homeostasis is therefore critical to unravelling their contributions to CNS disease and injury.

## Materials and methods

### Animals

Mice used for experiments were housed in specific pathogen-free facilities approved by the Association for Assessment and Accreditation of Laboratory Animal Care and procedures were performed in accordance with animal protocols approved by the Institutional Animal Care and Use Committee at The University of Colorado, Anschutz Medical Campus. Mouse lines used in this study include: (1) *Col1a1-GFP*: Tg(Col1a1-EGFP)#Dab (MGI 4458034; Yata et al., 2003); (2) *Col1a2-CreERT2*:Tg(Col1a2-cre/ERT,-ALPP)7Cpd (The Jackson Laboratory, #029567, RRID: IMSR_JAX:029567); (3) *Pdgfra-*EGFP: B6.129S4-*Pdgfratm11(EGFP)Sor*/J (The Jackson Laboratory, #0076699, RRID: ISMR_JAX:007669); (4) *Pdgfra-flox*: B6.Cg- *Pdgfra^tm8Sor^*/EiJ (The Jackson Laboratory, #006492, RRID: ISMR_JAX:006492). For immunohistochemistry and RNAscope *in situ* experiments in Figures 1 and 2, adult *Col1a1-GFP* or *Col1a2-CreERT2* mice were bred to produce P8 *Col1a1-GFP+* and *Col1a2-CreERT* (Cre negative) P21 pups. For cell culture experiments, adult *Col1a1-GFP* mice were bred to generate GFP+ and GFP- pups aged P3-4. For conditional knockout of PDGFRα, *Col1a2-CreERT2* mice heterozygous for Cre were crossed with *Pdgfra*-EGFP mice to generate offspring that were *Col1a2-CreERT2^Cre/+^*;*Pdgfra^EGFP/+^* . These offspring were then crossed with *Pdgfra* homozygous flox mice bred in with a tdTomato reporter allele (*Pdgfra^fl/fl^; Ai14^fl/fl^*) to generate *Col1a2- CreERT^Cre/+^*; *Ai14 ^fl/+^; Pdgfra^EGFP/fl^* cKO mutants and *Col1a2-CreERT^Cre/+^*; *Ai14 ^fl/+^; Pdgfra^+/flox^*conditional-heterozygous (cHet) littermate controls. Additional crosses of *Col1a2-CreERT2^Cre/+^*; *Pdgfra^EGFP/+^* and *Ai14^fl/fl^* mice were used to generate *Col1a2-CreERT2^Cre/+^*; *Ai14 ^fl/+^; Pdgfra^EGFP/+^* global-homozygous (gHet) and *Col1a2-CreERT2^Cre/+^*; *Ai14 ^fl/+^; Pdgfra^+/+^*wildtype controls. These pups were injected at P1, P2, and P3 with 50 μl of 1 mg/mL tamoxifen (Sigma-Aldrich, T5648) in corn oil (Sigma- Aldrich, C8267) to achieve recombination. We have previously shown that recombination of the *Ai14 ^fl/+^* tdTomato reporter does not occur in *Col1a2- CreERT2^Cre/+^* pups in absence of tamoxifen (Jones et al., 2023). Pups were collected at P10 for immunohistochemical analysis.

### Tissue collection, optical clearing, immunohistochemistry, and in situ RNAScope detection

For all immunohistochemistry and RNAscope *in situ* experiments, pups were humanely euthanized and transcardially perfused with 5mL PBS followed by 5mL 4% PFA. Brains were removed and post-fixed in 4% PFA at 4°C overnight, then incubated in 20% sucrose at 4°C for up to three days. For RNAscope *in situ* and protein immunohistochemistry in Figures 1, 2, and Supplemental Figure 1, brains were embedded and flash-frozen in TissueTek O.C.T (Sakura, Ref#: 4583) and sectioned coronally on a cryostat 12-15μm thick. For protein immunohistochemistry in Figures 3, 4 and Supplemental Figure 3, brains were embedded in 3% agarose (1:1 mixture of UltraPure LMP Agarose, Invitrogen, 16520-050, and Agarose, Fisher Bioreagents, BP1356) and sliced on a vibratome (Leica VT1200S) to a thickness of 100μm or 2 mm. For immunohistochemistry on cryosectioned and 100μm thick sections, brain sections were permeabilized in 0.1% Triton-X 100 for 10 minutes then blocked for 30 minutes in 2% lamb serum and 0.05% Triton-X 100 at room temperature. Slices were incubated in primary antibodies in block solution at 4°C overnight, followed by incubation in secondary antibodies in 0.05% Triton-X 100 for 1 hour at room temperature. For 2 mm-thick brain slices, our methods for clearing, staining, and imaging are described at length in <u>Jones et al., 2022.</u> The following antibodies were used: mouse anti-Collagen1a1 (1:100, Sigma-Aldrich, SAB1402151), goat anti- PDGFRα (1:100, R&D Systems, AF1062), rat anti-PDGFRβ (1:100, Novus, NBP1-43349), rat anti-CD31 (PECAM) (1:100, BD Pharmigen, 550274), rabbit anti-pan-laminin (1:100, Sigma Aldrich, NC1732938), rabbit anti-AQP4 (1:100, Alomone Labs, 249-323), goat anti-CD206 (1:100, R&D Systems, AF2535), with species-specific Alexa-Fluor (Invitrogen) secondary antibodies used at a dilution of 1:500. For the detection of PDGF receptor and ligand transcripts, the RNAScope Manual Detection Kit v.2.0 from ACDBio (Cat#: 323100) was used along with probes against *Pdgfra* (Cat#: 480661-C3)*, Pdgfrb* (Cat#: 411381-C3)*, Pdgfa* (Cat#: 411361)*, Pdgfb* (Cat#: 424651-C3), and *Pdgfc* (Cat#: 424651-C3). For vascular labeling in RNAScope tissue DyLight Tomato Lectin 649 (Vector Laboratories, DL-1178-1) was used.

### Image acquisition, quantification and statistical analysis

Images were obtained using Zeiss 780 and 900 LSM confocal microscopes and Zen Blue software (Zeiss) and processed in FIJI or IMARIS. Quantification and analysis of PVF coverage, cell depth and density in Figure 3 and Supplemental Figure 2 was conducted as previously described in Jones et al., 2023 using FIJI. Briefly, individual vessels in which the entire length of vessel could be observed and measured were first selected from maximum z-projected images with only the labeled vessel channel (lectin) shown. For each vessel the distance between the meninges and the furthest traveled PVF, and the distances between the first 5 PVFs were measured to obtain cortical depth measurements (Sup. Fig 2A, Figure 3F). Percent coverage of individual vessels was obtained by dividing the cortical depth of the furthest traveled PVF by the total vessel length (Figure 3C). Density of PVFs on a vessel was calculated by dividing the cortical depth of the furthest traveled PVF by the number of PVFs contained within that vessel length and multiplying by 100 μm to normalize measurements (Sup. Fig 2B). For quantification of density of meningeal fibroblasts in Supplemental Figure 1B, the total numbers of DAPI+/tdTomato+ cells were counted in the meninges using the Cell Counter plugin in FIJI and divided by the length of the meninges measured (in μm) in the field-of-view, then multiplied by 100 μm to normalize. All statistical analyses were performed in GraphPad Prism (v.10.4.1). Normality tests were performed before statistical analyses where necessary. Respective statistical tests and replicate numbers are reported in figure legends. Where shown, lines and error bars represent mean and standard deviation.

## Supporting information

Supplemental Figure 1, 2 Graphical Abstract

## Acknowledgments

The authors would like to thank the members of the Siegenthaler Lab and members of HEJ’s thesis committee (Dr. Santos Franco, Dr. Linda Barlow, Dr. Fabrice Dabertrand and Dr. Joe Brzezinski) for fruitful discussions and feedback that helped shape this project. This work was supported by funding from the NIH, specifically NINDS and NICHD (F31NS125875 and T32HD007186 to HEJ, R01 NS098273 and NS131823 to JAS). All schematics in the figures were made with BioRender.

## Author Contributions

J.S. and H.E.J conceived the research. J.S. and K.F. supervised the research. J.S., H.E.J. and K.A. designed the experiments. H.E.J and K.A. carried out the experiments. K.F. provided essential mouse lines.

