## Supplemental Figure 1, 2 Graphical Abstract for "“PDGFRα is required for postnatal cerebral perivascular fibroblast development”"

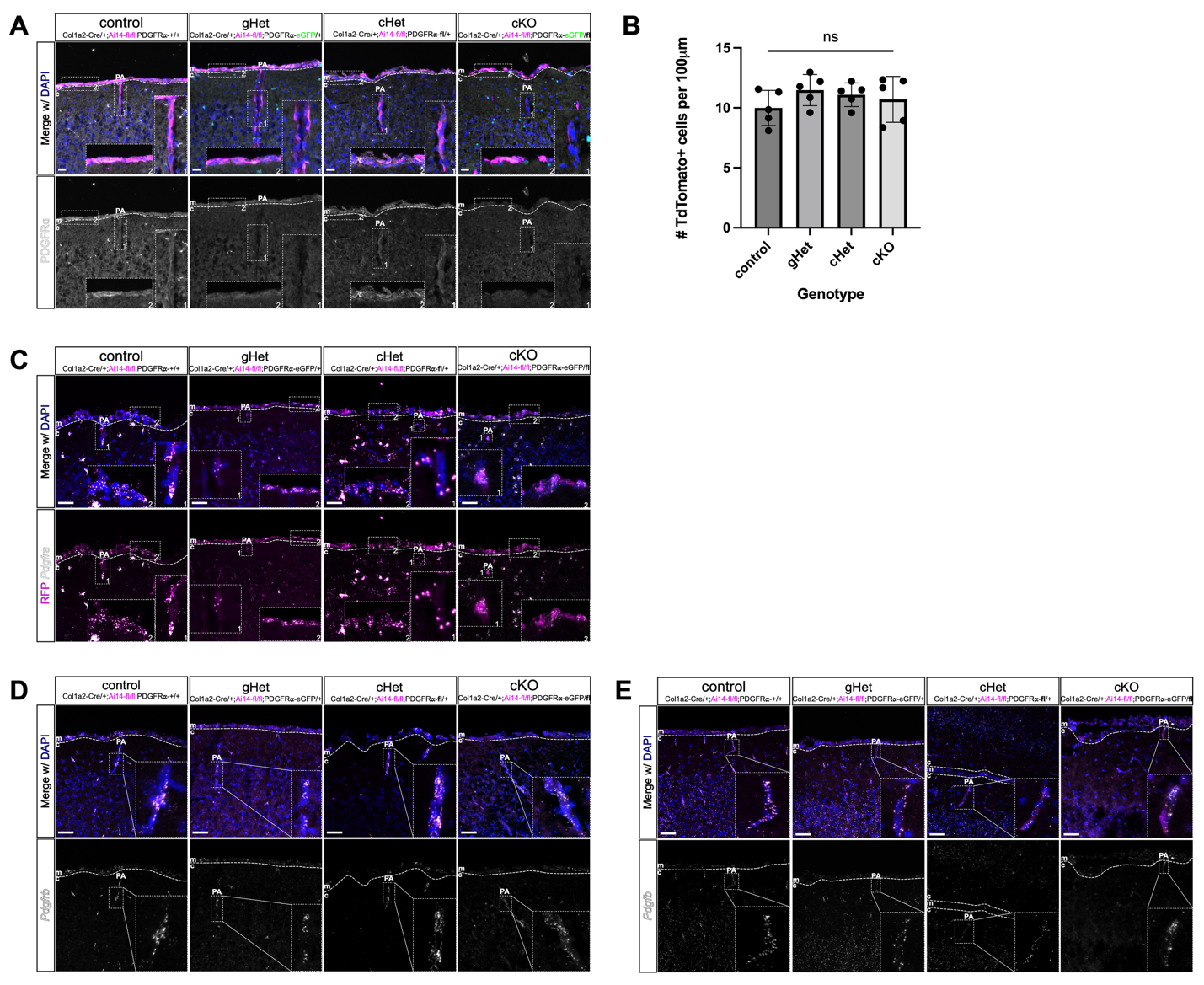

**Supplemental Fig 1: Validation of genetic conditional knockout of PDGFRα from PVFs *in vivo*** (**A**) Confocal images of meninges and penetrating vessels in P10 control, cHet, gHet and cKO mice showing fibroblasts (RFP, magenta), eGFP (green, gHet and cKO only), PDGFRα protein staining (white). (**B**) Graph showing density of meningeal fibroblasts represented as the number of TdTomato+ cells counted per 100μm length of meninges, N = 1 biological replicate per genotype, individual points represent individual fields-of-view counted (5 per animal). One-way ANOVA with multiple comparisons reveals no significant differences between genotypes. (**C**) Confocal images of meninges and penetrating vessels in P10 control, cHet, gHet and cKO mice showing fibroblasts (RFP, magenta) and RNAscope detection of *Pdgfra* (white). (**D**-**E**) Confocal images of meninges and penetrating vessels in P10 control, cHet, gHet, and cKO mice showing fibroblasts (RFP, magenta) and RNAscope detection of (**D**) *Pdgfrb* or (**E**) *Pdgfb*. Scale bars (**A**) 25μm, (**C, D, E**) 50μm. **m**: meninges, **c**: cortex, **PA**: penetrating arteriole; dashed line denotes border between meninges & cortex, dashed boxes indicate areas of inset.

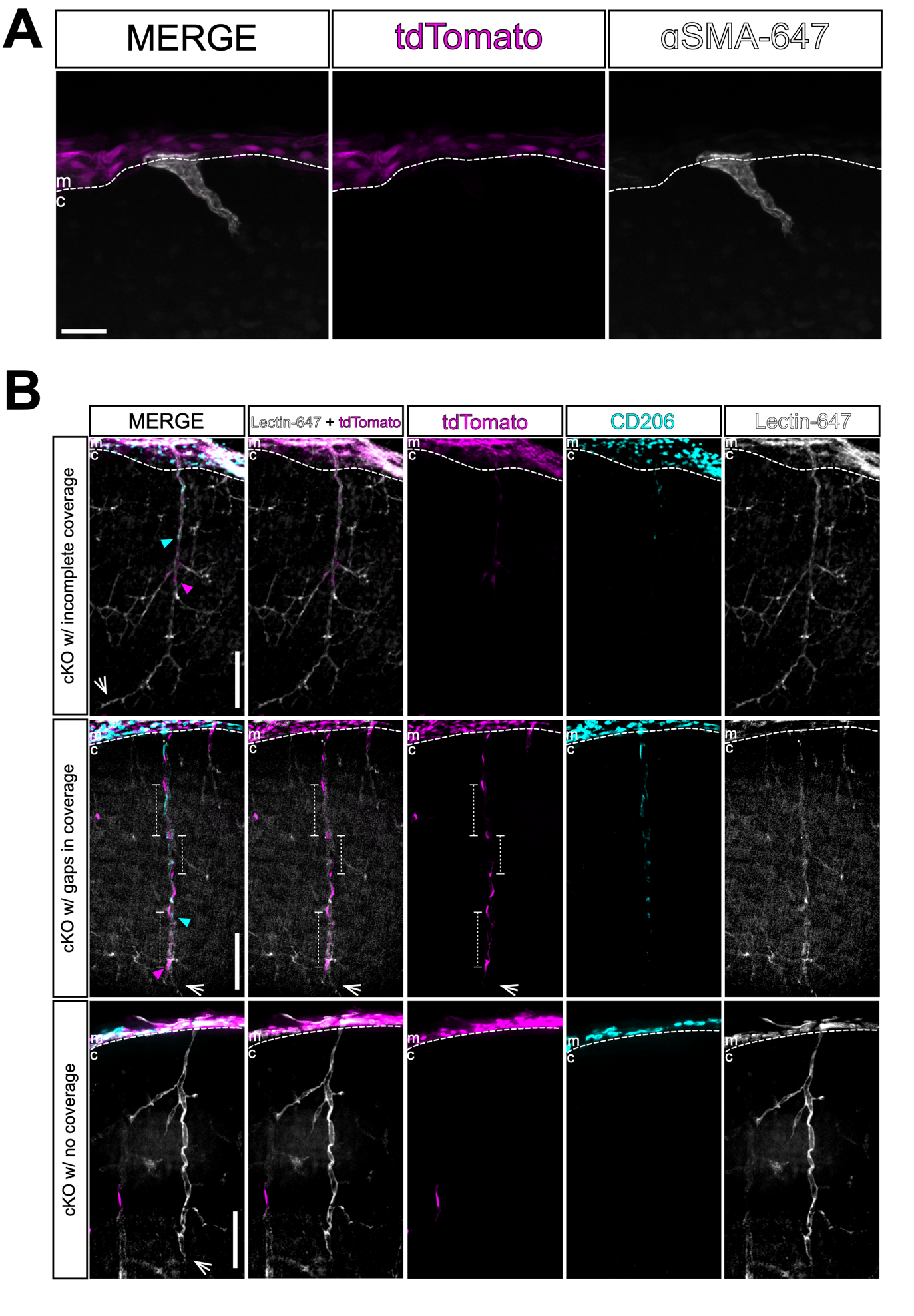

**Supplemental Figure 2: Characterization of PVS components on PDGFRα cKO vessels.**

**(A**) ) Confocal image of meninges and penetrating in cKO mutant at P10 that lacks PVFs, showing meningeal fibroblasts labeled with tdTomato (magenta), with vSMCs labeled by αSMA (white). (**B**) Confocal images of meninges and penetrating vessels with varying levels of PVF coverage in cKO mutant at P10 showing fibroblasts labeled with tdTomato (magenta), PVMs labeled by CD206 (cyan), and vessels labeled with lectin (white). **m**: meninges, **c**: cortex; dashed line denotes border between meninges and cortex. Scale bars (**A**) 25μm, (**B**) 100μm.

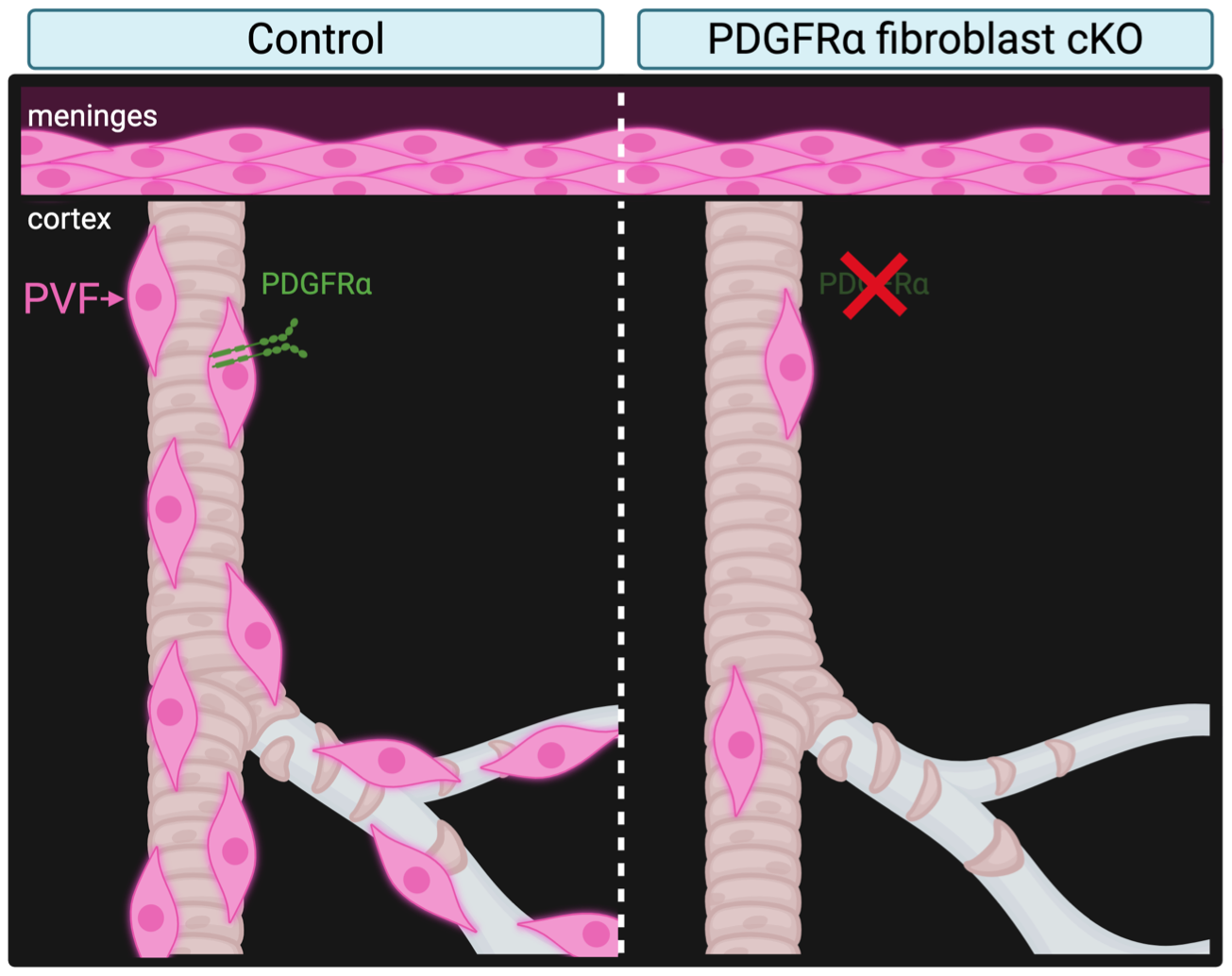

**Graphical Abstract.** Conditional deletion of PDGFRα from CNS fibroblasts disrupts cerebrovascular coverage by PVFs in the postnatal mouse brain.
